# The E3 Ubiquitin Ligase RNF5 Facilitates SARS-CoV-2 Membrane Protein-Mediated Virion Release

**DOI:** 10.1101/2021.02.28.433287

**Authors:** Zhen Yuan, Bing Hu, Yulei Wang, Xuan Tan, Hurong Xiao, Mengzhen Yue, Kun Cai, Ke Tang, Binbin Ding

## Abstract

As enveloped virus, SARS-CoV-2 membrane protein (M) mediates viral release from cellular membranes, but the molecular mechanisms of SARS-CoV-2 virions release remain poorly understood. Here, we performed RNAi screening and identified the E3 ligase RNF5 which mediates ubiquitination of SARS-CoV-2 M at residue K15 to enhance the interaction of viral envelope (E) with M. M-E complex ensures the uniform size of viral particles for viral maturation and mediates viral release. Moreover, overexpression of M induces complete autophagy which is dependent on RNF5-mediated ubiquitin modification. M inhibits the activity of lysosome protease, and uses autolysosomes for virion release. Consequently, all these results demonstrate that RNF5 mediates ubiquitin modification of SARS-CoV-2 M to stabilize the M-E complex and induce autophagy for virion release.

## MAIN TEXT

## Introduction

SARS-CoV-2 belongs to *betacoronavirus* of the family Coronaviridae which are enveloped viruses with a single-strand, positive-sense RNA genome (1, 2). Its diameter is about 65-125 nm. SARS-CoV-2 consists of four major structural proteins: spike (S), envelope (E), membrane (M) and nucleocapsid (N). Similar with other RNA viruses, the genomic RNA replication, mRNA transcription and proteins synthesis of coronavirus occur in cytoplasm (3). The newly synthesis structure proteins and the RNA genome are assembled into virions, and M drives the virions bud into the lumen of the endoplasmic reticulum-Golgi intermediary compartment (ERGIC) for the further modification and maturation (3, 4). M oligomerization of mouse hepatitis virus (MHV) mediated by its transmembrane domain is believed to allow the formation of a lattice of M (5). S and E are integrated into the lattice through interacting with M (6). Different from MHV that E and M proteins co-expressed in cells are necessary and sufficient for VLPs release, M/E/N of SARS-CoV are all required for efficient assembly and release, but the mechanisms are poorly understood (7). Virions release is essential step in the release of the enveloped virus particles that ultimately separates virion and host membranes. VLP (virus-like particle) systems could be used to determine the individual roles of different viral and cellular proteins in virions release and could be used for vaccine development. Therefore, by using this convenient method to screen and identify the key cellular proteins functioned in virion release, we can provide new insight into the details of viral maturation and advance our understanding of the virus-cells interaction.

Ubiquitin modification is important for virus replication. Ubiquitin is enriched in retrovirus particles and a variable fraction of the major retroviral structural protein (Gag) can be ubiquitinated (8, 9). Additionally, ubiquitination of cellular transmembrane proteins can signal the recruitment of class E machinery (VPS), a popular model is that deposition of ubiquitin on viral structural proteins mediates class E machinery recruitment (10). Viral M generally contains core consensus amino acid motifs called late (L) domains, which are essential for efficient viral release (11). Studies on parainfluenza virus found that M-mediated VLP production by ubiquitination (12, 13), proteasome inhibitor treatments block the release of PIV5, NiV, and SeV (14–16). Potential ubiquitination of MeV M has observed in cells and ESCRT factors such as ALIX can bind to ubiquitin and enhance viral release (17, 18). So far, an increasing number of studies have identified the E3 ligase in host responses to virus infection and ubiquitinated viral proteins to regulate virus replication. TRIM6 ubiquitinates the Ebola Virus VP35 to promote replication (19), TRIM69 restricts dengue virus (DENV) replication by ubiquitinating viral NS3 (20), TRIM26 ubiquitinates HCV NS5B to promote the NS5B-NS5A interaction for replication (21), MARCH8 ubiquitinates the HCV NSP2 and mediates viral envelopment (22), RNF 121 is a potent regulator of adeno-associated viral genome transcription (23). However, whether ubiquitination is required for SARS-CoV-2 release, and what are the E3 ligase and the deubiquitinating enzyme have not been determined.

RING finger protein family (RNF) have been demonstrated in the regulation of antiviral responses. RNF128 promotes innate antiviral immunity through K63-linked ubiquitination of TBK1 (24), RNF90 negatively regulates cellular antiviral responses by targeting MITA for degradation (25), RNF11 targets TBK1/IKKi kinases to inhibit antiviral signaling (26). RNF5, also known as RMA1, is a RING finger protein and a membrane-anchored (ER and/or mitochondria) E3 ubiquitin ligase, which anchored to the ER membrane via a single transmembrane, spanning the domain located within the C-terminal region. It is implicated in ERAD, cell motility and also negative regulation of autophagy and ER stress (27–30). Several studies have shown the connection between RNF5 and antiviral response: RNF5 negatively regulates virus-triggered signaling by targeting STING and MAVS for ubiquitination and degradation at mitochondria (31, 32); Newcastle Disease Virus V Protein degrades MAVS by recruiting RNF5 to polyubiquitinate MAVS (33). So far, we have limited knowledge about whether RNF5 could function as the E3 ligase of viral proteins to regulate viral release.

In this study, we identified RNF5 as the E3 ligase of SARS-CoV-2 M by using RNAi screening. A mechanistic study demonstrated that RNF5 regulates virion release by enhancing the interaction of M with E. Furthermore, we showed that M induces autophagy which is dependent on RNF5-mediated ubiquitin modification. We also found that M inhibits the activity of lysosome protease to block the degradation of autolysosomes, and uses autolysosomes for egress. All in all, our findings draw out the formation and regulation mechanisms of SARS-CoV-2 VLPs and provide molecular details of SARS-CoV-2 virion release. We identify RNF5 as the E3 ligase for ubiquitination of M which will be helpful in the development of novel therapeutic approaches.

## Results

### RNF5 Promotes SARS-CoV-2 Virion Release

We began investigating the mechanisms of SARS-CoV-2 assembly and release by using VLPs, as VLPs systems had been proved to be useful tools for studying the viral assembly and release processes of many enveloped viruses. We first transiently expressed M alone, M/E and M/N of SARS-CoV-2, and culture medium were collected and subjected to ultracentrifugation to pellet VLPs. We found that only M/E co-expression resulted in a significant VLPs formation, not M alone (**Fig. 1A**). To further confirm that the pellet VLPs were indeed the membrane-bound VLPs, we then treated VLPs with trypsin or/and Triton X-100. No significant digestion of M was observed either in trypsin or Triton X-100, in contrast, under trypsin plus Triton X-100 treatment, M was completely degraded (**Fig. 1B**). These data suggested that SARS-CoV-2 E is required for M mediated VLPs release. Then we used this convenient assay to investigate the mechanism(s) of SARS-CoV-2 release. To identify host factors essential for virion release, we performed a small scale of RNAi screening targeting candidates which are on the list from SARS-CoV-2 M IP/MS (34), and found that knockdown of RNF5 (Ring Finger Protein 5), an ER-localized E3 ligase, significantly reduced virion release (**Fig. S1A**). We confirmed the interaction of M with RNF5 via co-IP: endogenous RNF5 coimmunoprecipitated with M (**Fig. 1C**). RNF5 colocalized well with M in cytoplasm (**Fig. 1D**). TMD of RNF5 and CTD of M were critical for M-RNF5 interaction (**Fig. S1B and S1C**). Next, we confirmed that knockdown of RNF5 did decrease the virion release (**Fig. 1E**). Over-expression of RNF5 significantly increased VLPs release while its catalytic dead mutant RNF5-C42S reduced VLPs release may be due to the domain negative effect (**Fig. 1F**), and RNF5-C42S still interacted with M (**Fig. S1D**), suggesting that RNF5 promotes virion release which is dependent on its E3 ligase activity. Furthermore, to exclude potential off-target effects of siRNA, we performed rescue experiments in *rnf5* KO cells and found that wild-type RNF5, but not C42S rescued the reduction of VLPs release (**Fig. 1G**). Next, we asked whether RNF5 could modulate the maturation and release of SARS-CoV-2. We infected wild type and RNF5 KD Vero cells with SARS-CoV-2 and performed plaque assay. The extracellular viral production was lower in RNF5 KD cells than wild type cells (**Fig. 1H and S1E**). To exclude the possibility of infectibility defect in RNF5 KD cells, we evaluated the viral gene expression in extracellular and intracellular via real-time PCR and found that intracellular SARS-CoV-2 ORF1ab gene expression was slightly decreased while extracellular ORF1ab expression was significantly decreased in RNF5 KD cells than wild type cells (**Fig. 1I**). Taken together, our results suggested that RNF5 facilitates SARS-CoV-2 release.

**Figure 1.**
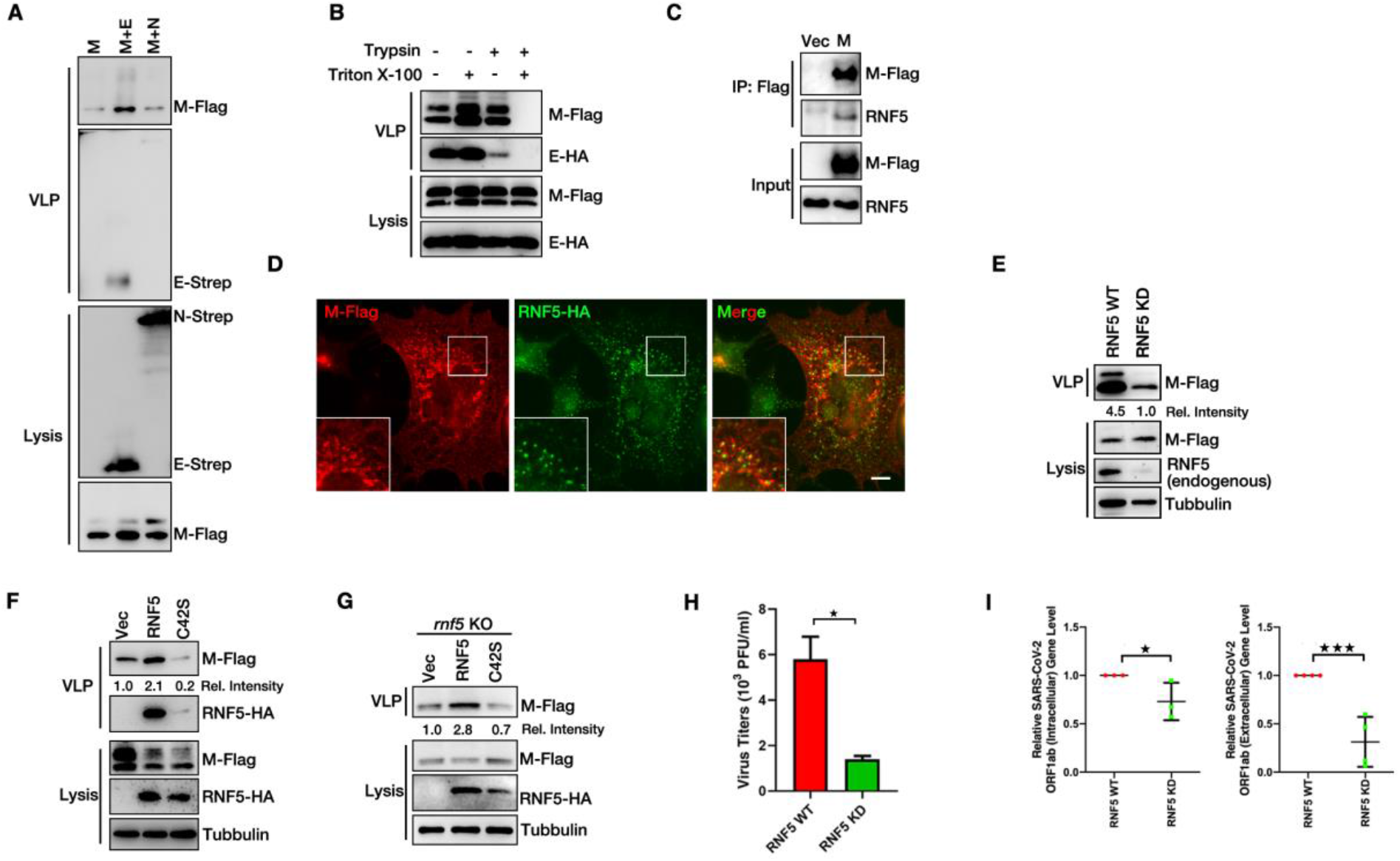
RNF5 Regulates SARS-CoV-2 Viral Release. (A) HEK293T cells were transfected with the indicated plasmids for 36 h, and the VLP release assay was performed as described in Materials and Methods and analyzed via western blotting (WB). (B) HEK293T cells were transfected with SARS-CoV-2 M-Flag and E-HA for 36 hr. A protease protection assay of VLPs was performed as described Experimental Procedures and then analyzed via WB. (C) HEK293T cells were transfected with M-Flag for 36 h. Lysates were subjected to Flag IP and analyzed via WB. (D) AD293 cells were transfected with M-Flag and RNF5-HA for 36h, and cells were analyzed via immunofluorescence. Flag antibody was used for tracking M and HA antibody was used for tracking RNF5. Scale bar: 10 μm. (E) RNF5 WT and KD Vero-E6 cells were transfected with M-Flag and E-HA for 36h. Lysates and corresponding purified VLPs were analyzed via WB. (F) HEK293T cells were transfected with M-Flag, E-HA and RNF5-HA or C42S-HA for 36 h. Lysates and corresponding purified VLPs were analyzed via WB. (G) *rnf5* KO HEK293T cells were transfected with M-Flag, E-HA and RNF5-HA or C42S-HA for 36 h. Lysates and corresponding purified VLPs were analyzed via WB. (H) RNF5 WT and KD Vero-E6 cells were infected with SARS-CoV-2 (WBP-1) for 24 h and then mediums were collected and analyzed via Plaque Assay as described in Materials and Methods. (I) The level of SARS-CoV-2 ORF1ab mRNA of intracellular and extracellular from (H) were measured via real-time PCR. Error bars, mean ± SD of three experiments (n = 3). Student’s t test; *p < 0.05; ***p < 0.001.

### SARS-CoV-2 M Interacts with E which is Required for Virion Release

Then we asked how RNF5 promotes virion release. We have shown that M and E are both required for VLPs release and RNF5 interacts with M, then we speculated that RNF5 promotes virion release by targeting M. So, we first sought to determine the mechanism(s) of how M-E mediate virion release. We used NanoSight NS300 (Malvern) to analyze the size distribution of VLPs and found that the size of VLPs from M alone were inhomogeneous with the diameters from 40nm to 600 nm and showed several peaks, suggesting M alone fails to efficiently release as complete VLPs, and the size of VLPs from M-E co-expression were uniform with average diameter 144 nm (**Fig. 2A**), suggesting that E interacts with M to ensure the uniform size of viral particles. The kinetics of E expression parallels the increasement of M-mediated VLPs formation (**Fig. 2B**). M self-interaction was required for VLPs formation, as mutant M_ΔCTD_ failed to interact with M (**Fig. 2C**) and lost the ability to release as VLPs (**Fig. 2D**). E expression parallels the increasement of M self-interaction in co-immunoprecipitation assay (**Fig. 2E**). In the presence of the crosslinker disuccinimidyl suberate (DSS), we found that M existed in monomeric, dimeric and oligomeric states, and E expression enhanced the homo-oligomerization of M (**Fig. 2F**). These results demonstrated a critical role of SARS-CoV2 E in viral M homo-oligomerization and effective virion release. Similar with SARS-CoV, M interacted with E (**Fig. 2G**). C-terminal of M was required for its interaction with E, as mutant M_ΔCTD_ failed to bind to E (**Fig. 2G**) and release as VLPs (**Fig. 2D**), and CTD of M alone was sufficient to interact with E (**Fig. 2H**), suggesting that M interacts with E via its CTD and this interaction is critical for VLPs release. M_ΔNTD_ showed less interaction with E (**Fig. 2G**) and VLP formation (**Fig. 2D**). Deletion of either one of three TMDs (transmembrane domain) of M had little impact on M self-interaction, M-E interaction and VLPs release (**Fig. 2C, 2G and 2I**). Remarkably, M_ΔCTD_, TMD deleted mutant (Δ20-100aa) and CTD plus NTD deleted mutant (20-100aa) completely lost their ability to form VLPs (**Fig. 2D**). Thus, M uses its CTD for M-M and M-E interaction which are critical for VLPs release.

**Figure 2.**
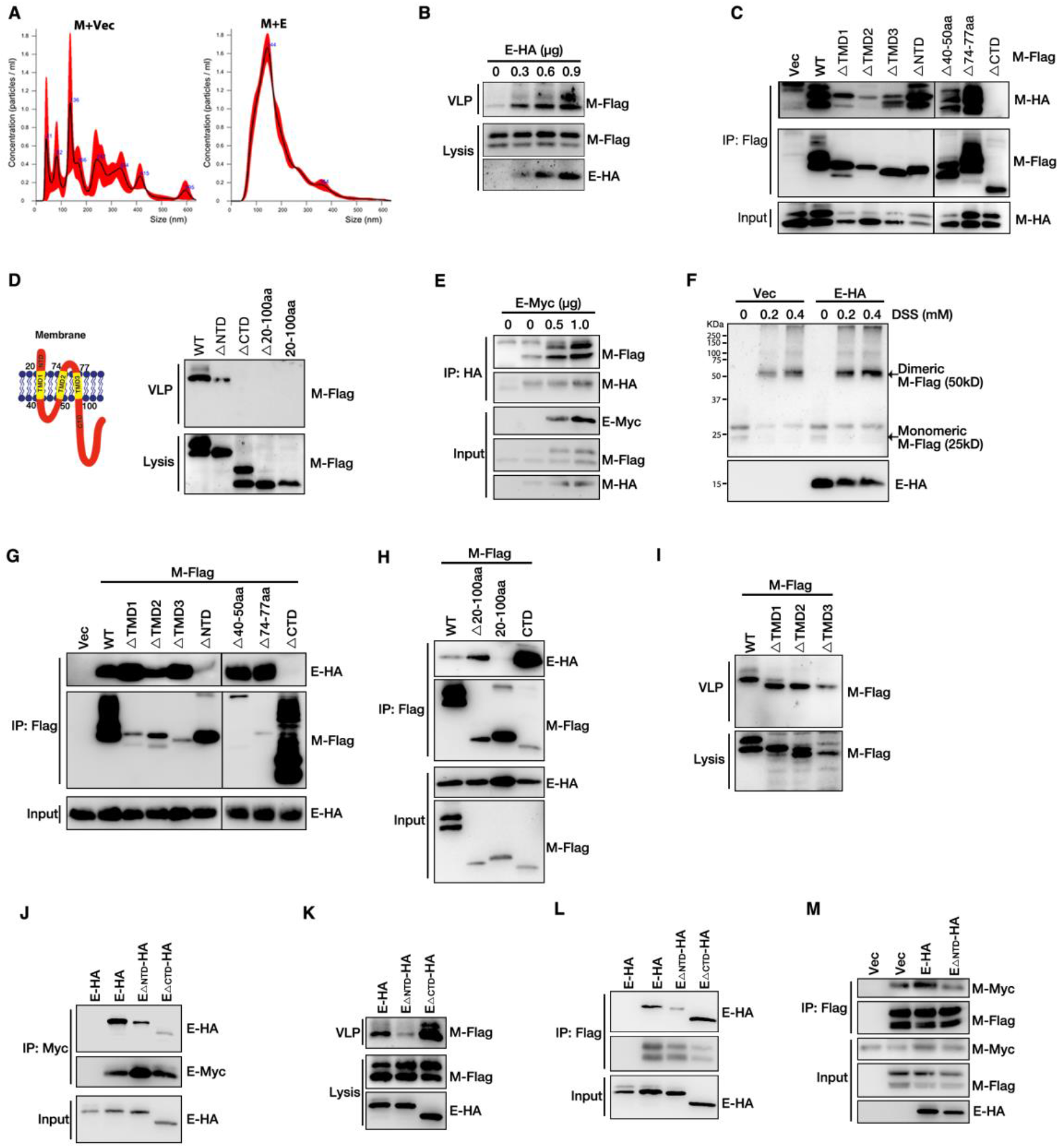
SARS-CoV-2 M Interacts with E to Mediate Viral Release. (A) HEK293T cells were transfected with M-Flag with or without E-HA for 36 h, then size of purified VLPs were analyzed via NanoSight NS300 (Malvern). (B) HEK293T cells were transfected as indicated for 36 h, and the VLP release assay was performed and then analyzed via WB. (C) HEK293T cells transfected with M-HA and M-Flag or its mutants for 36 h, and subjected to Flag IP and analyzed via WB. (D) Schematic diagrams of wild-type M. HEK293T cells were transfected with plasmids as indicated for 36 h. Lysates and corresponding purified VLPs were analyzed via WB. (E) HEK293T cells were transfected with M-HA and M-Flag and E-Myc for 36 h, and subjected to HA IP and analyzed via WB. (F) HEK293T cells were transfected with M-Flag and E-HA for 36 h, then cells were cross-linking by treating with DSS for 30 min and then lysates were analyzed via WB. (G) HEK293T cells were transfected with E-HA and M-Flag or its mutants for 36 h, and subjected to Flag IP and analyzed via WB. (H) HEK293T cells were transfected with E-HA and M-Flag or its mutants for 36 h, and subjected to Flag IP and analyzed via WB. (I) HEK293T cells were transfected with E-HA and M-Flag or M deleted TMD1, TMD2 or TMD3 for 36 h, and the VLP budding assay was performed and analyzed via WB. (J) HEK293T cells were transfected with E-Myc and E-HA or its mutants for 36 h, and subjected to Myc IP and analyzed via WB. (K) HEK293T cells were transfected with M-Flag, E-HA and mutants for 36 h. Lysates and corresponding purified VLPs were analyzed via WB. (L) HEK293T cells were transfected with M-Flag and E-HA or its mutants for 36 h, and subjected to Flag IP and analyzed via WB. (M) HEK293T cells were transfected with M-Flag, M-Myc, E-HA and mutant for 36 h. Lysates were subjected to Flag IP and then analyzed via WB.

Previous study showed that SARS-CoV E forms ion channel on membrane via homo-oligomerization (35). Thus, we sought to determine whether SARS-CoV-2 E could exist homo-oligomerization and this oligomerization plays an important role in VLPs release. We generated several truncations and found that E_ΔCTD_ showed less self-interaction in a co-IP assay (**Fig. 2J**). To our surprise, E_ΔCTD_ still promoted VLPs release (**Fig. 2K**), suggesting that E homo-oligomerization is not required for VLPs release. Remarkably, E_ΔNTD_ mutant failed to interact with M (**Fig. 2L**) and lost its ability to promote VLPs release (**Fig. 2K**). Furthermore, we found that E_ΔNTD_ failed to enhance the self-interaction of M (**Fig. 2M**). Thus, E-M interaction, not E homo-oligomerization is essential for VLPs release. Taken together, these data suggested that E interacts with M to enhance the self-interaction of M, which ensures the uniform size of viral particles and thus promotes virion release.

### RNF5 Promotes M-E Interaction

Next, we want to determine whether RNF5 regulates M-E interaction. Over-expression of RNF5 had little impact on protein level of M (**Fig. 3A**). Wild-type RNF5, but not C42S mutant increased the M-E interaction (**Fig. 3B**) while had no effect on M self-interaction (**Fig. 3C**), suggesting that RNF5 enhances the interaction of M with E which is dependent on its E3 ligase activity. Similar result was obtained in rescue experiments in *rnf5* KO cells: wild-type RNF5, but not C42S rescued the M-E interaction (**Fig. 3D**). We further used NanoSight NS300 (Malvern) to analyze the size distribution of VLPs from RNF5 KD cells and found that knockdown of RNF5 lead to inhomogeneous VLPs, suggesting that RNF5 ensures the uniform size of viral particles (**Fig. 3E**). Taken together, these results suggested that RNF5 enhances the interaction of M with E to promote virion release and ensure the uniform size of viral particles.

**Figure 3.**
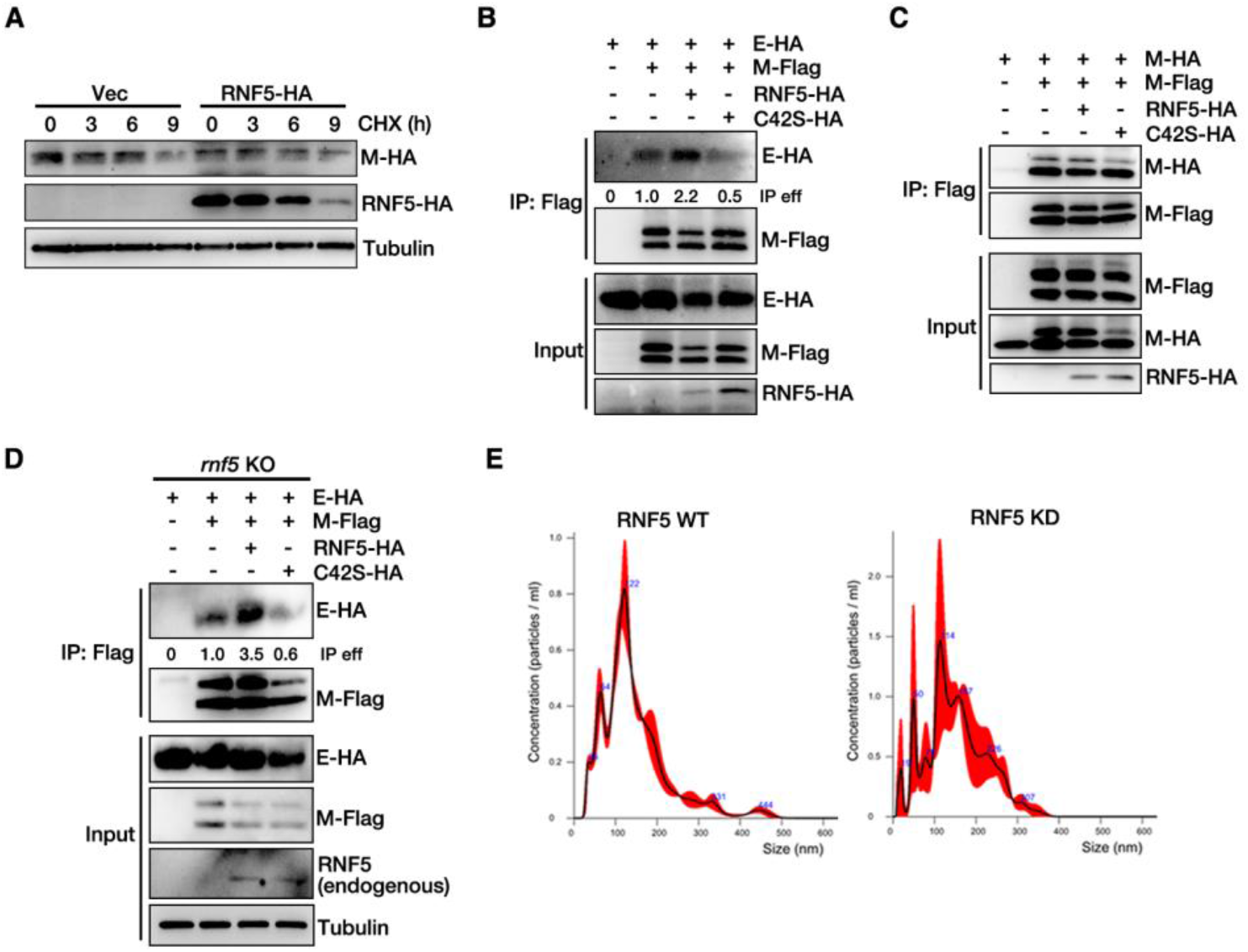
RNF5 Promotes the Interaction of M with E for Viral Release. (A) M-HA expression stable HEK293T cells were transfected with or without RNF5-HA and treated with CHX for indicated hours, and cells lysis were analyzed via WB. (B) HEK293T cells were transfected with M-Flag, E-HA and RNF5-HA or mutant C42S-HA for 36 h. Lysates were subjected to Flag IP and analyzed via WB. (C) HEK293T cells were transfected with indicated plasmids for 36 h, and subjected to Flag IP and analyzed via WB. (D) *rnf5* KO HEK293T cells were transfected with indicated plasmids for 36 h, and subjected to Flag IP and analyzed via WB. (E) *rnf5* KO HEK293T cells were transfected with M-Flag and E-HA with or without RNF5-HA for 36 h, then size of purified VLPs were analyzed via NanoSight NS300 (Malvern).

### Ubiquitination of SARS-CoV-2 M Mediated by RNF5 is Critical for Efficient Virion Release

RNF5 is an E3 ligase and enhances the interaction of M with E which is dependent on its E3 ligase activity. Thus, we speculated that RNF5 ubiquitinates M to regulate M-E interaction and virion release. We first confirmed that M can be ubiquitinated (**Fig. 4A**). Overexpression of wild-type RNF5, but not RNF5-C42S, enhanced ubiquitination level of M (**Fig. 4B**). We also performed rescue experiments in *rnf5* KO cells and found that knockout of *rnf5* completely abolished ubiquitination of M, and wild-type RNF5, but not C42S rescued ubiquitination of M (**Fig. 4C**). Overexpression of RNF5 only increased the K63-linked polyubiquitination and had minor effect on the K48-linked polyubiquitination of M (**Fig. 4D**). These results suggested that RNF5 acts as the E3 ligase for K63-linked ubiquitination of M. We next sought to define the critical lysine residues of ubiquitin modification in M which may be responsible for VLPs release. Deletion of TMD (Δ20-100aa) in M had no effect on ubiquitin modification, and CTD plus NTD deleted mutant (20-100aa) showed less ubiquitination (**Fig. 4E**), suggesting that ubiquitin modification motif is on non-transmembrane domain. We further found that M_ΔNTD_ completely lost its ability for ubiquitination (**Fig. 4F**). Two lysine repeats K14/K15 are localized to the N terminus and four lysine repeats K162/K166/K180/K205 are localized to the C terminus of M (**Fig. 4G**). We found that only residue K15 was mainly responsible for M ubiquitination, as K15R mutant completely abolished the ubiquitin modification (**Fig. 4H**). These results indicate that residue K15 is a main ubiquitin modification site in M. Ubiquitination modification can result in altering its localization or binding partners. Next, we sought to determine whether ubiquitination in M plays potential roles in M-E interaction and SARS-CoV-2 virion release. To verify this assumption, we performed co-IP and VLP assay. Notably, only K15R mutant showed less M-E interaction and dramatic reduction of VLPs release (**Fig. 4I**). These data suggested that RNF5 ubiquitinates M at the residue of K15 to enhance the interaction of M and E.

**Figure 4.**
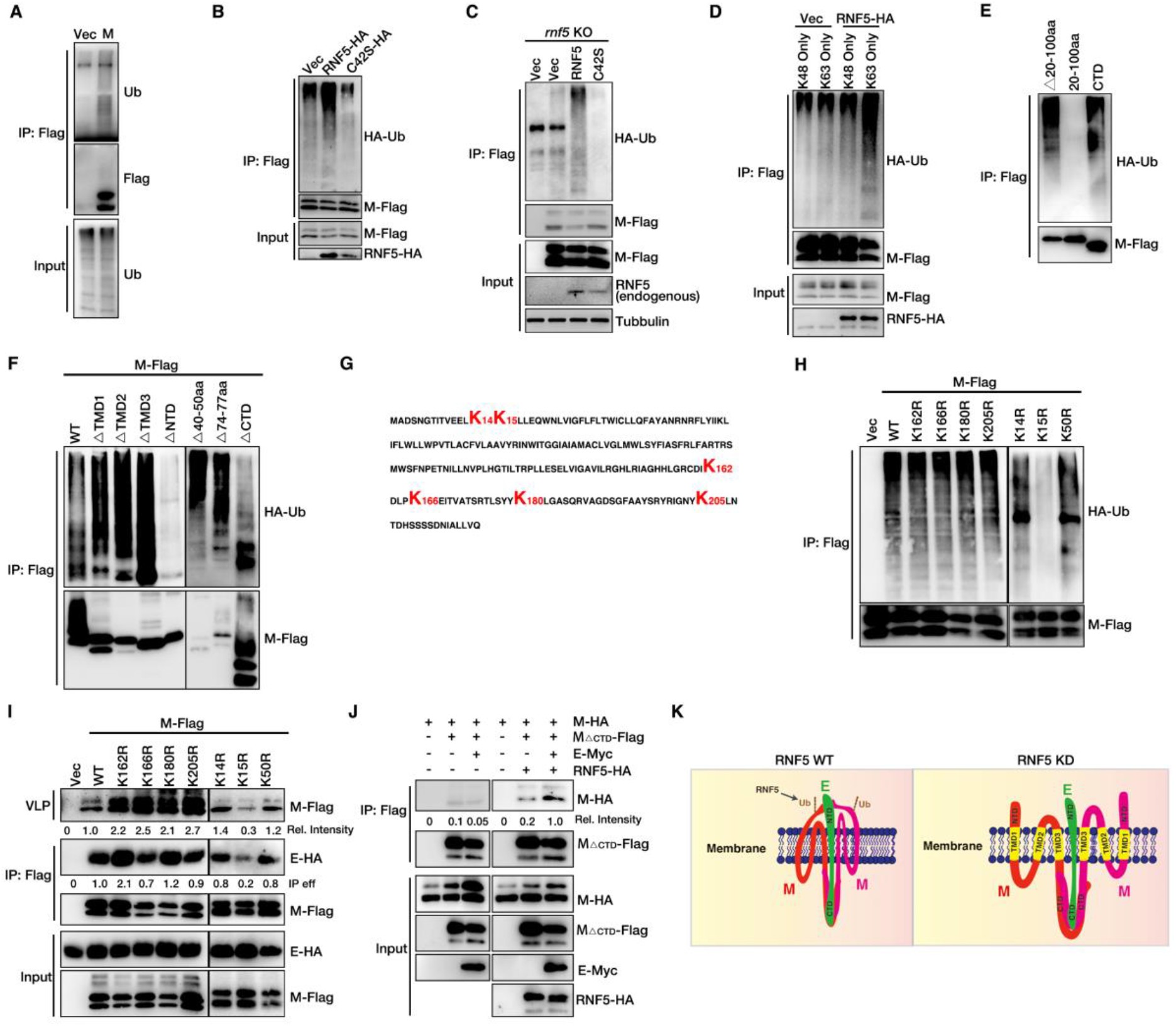
Ubiquitination of SARS-CoV-2 M Mediated by RNF5 is Critical for Viral Release. (A) HEK293T cells were transfected with HA-Ub and M-Flag for 36 h, and subjected to Flag IP and analyzed via WB. (B) HEK293T cells were transfected with M-Flag, HA-Ub and RNF5-HA or mutant C42S-HA for 36 h. Lysates were subjected to Flag IP and analyzed via WB. (C) *rnf5* KO HEK293T cells were transfected with M-Flag, HA-Ub and RNF5-HA or C42S-HA for 36 h. Lysates were subjected to Flag IP and analyzed via WB. (D) HEK293T cells were transfected M-Flag with HA-UbK48 Only or HA-UbK63 Only with or without RNF5-HA for 36 h. Lysates were subjected to Flag IP and analyzed via WB. (E) HEK293T cells were transfected M-Flag or mutants with HA-Ub. Lysates were subjected to Flag IP and analyzed via WB. (F) HEK293T cells were transfected with HA-Ub and M-Flag or its mutant for 36 h, and subjected to Flag IP and analyzed via WB. (G) Amino acid sequence of SARS-CoV-2 M. (H) HEK293T cells were transfected M-Flag or mutants with HA-Ub. Lysates were subjected to Flag IP and analyzed via WB. (I) HEK293T cells were transfected with E-HA, M-Flag and mutants for 36 h. Lysates were subjected to Flag IP and corresponding purified VLPs analyzed via WB. (J) HEK293T cells were transfected with indicated plasmids for 36 h, and subjected to Flag IP and analyzed via WB. (K) Model of RNF5 regulates M-E interaction.

We had shown that: 1) M only used its CTD for self-interaction and M-E interaction; 2) E helped to mediate the oligomerization of M; 3) Ubiquitin modification site was on the NTD of M and critical for both M-E interaction and VLPs release. Based on these results, we speculated that RNF5 ubiquitinates M at the residue of K15 to enhance the interaction of NTD of M with E, thus increases the stability of M-E complex on membrane to ensure the uniform size of VLPs and viral maturation. To verify this hypothesis, we tested the self-interaction of M_ΔCTD_ with or without E and RNF5 via co-IP. The result suggested that M_ΔCTD_ alone showed less self-interaction, expression of E and RNF5 significantly increased the self-interaction of M_ΔCTD_ (**Fig. 4J**). Taken together, these results indicated that E binds to the CTD of M and promotes the oligomerization of M, RNF5 ubiquitinates M at the residue of K15 to enhance the interaction of M NTD with its self and E, thus enhancing the stability of M-E complex to promote VLPs assembly and release (**Fig. 4K**).

### SARS-CoV-2 M Induces Autophagy and Inhibits Lysosome Protease for Virion Release

Our previous study indicated that M of HPIV3 colocalized with LC3 and autophagosome enhances the ability of virions binding to membrane and release (36). A recent study showed that SARS-CoV-2 uses lysosomes for egress instead of the biosynthetic secretory pathway via lysosome deacidification, inactivation of lysosomal degradation enzymes (37). Then we asked whether SARS-CoV-2 hijacks autophagosome/autolysosome for release. We first treated cells with Kifunensine (inhibitor of ER-associated degradation, ERAD) or Monensin (inhibitor of Golgi mediated transport) or Brefeldin A (inhibit the secretion targeting Golgi complex disassembles and redistributes into ER) or CQ (inhibitor of lysosome degradation) or Torin1 (inhibitor of mTORC1, inducer of autophagy) and subjected to VLPs assay. We found that Monensin treatment slightly decreased VLPs release and Kifunensine or Brefeldin A treatment had no effect on VLPs release (**Fig. 5A**), suggesting that virion release is not dependent on ERAD and ER-Golgi trafficking. Remarkably, CQ or Torin1 treatment significantly enhanced VLPs release (**Fig. 5B**), and knockdown of Atg7, the key gene for autophagy induction, significantly reduced the VLPs release (**Fig. 5C**), and LC3-I and LC3-II were co-released with M mediated VLPs (**Fig. 5D**), suggesting that M uses autophagosome/autolysosome for release.

**Figure 5.**
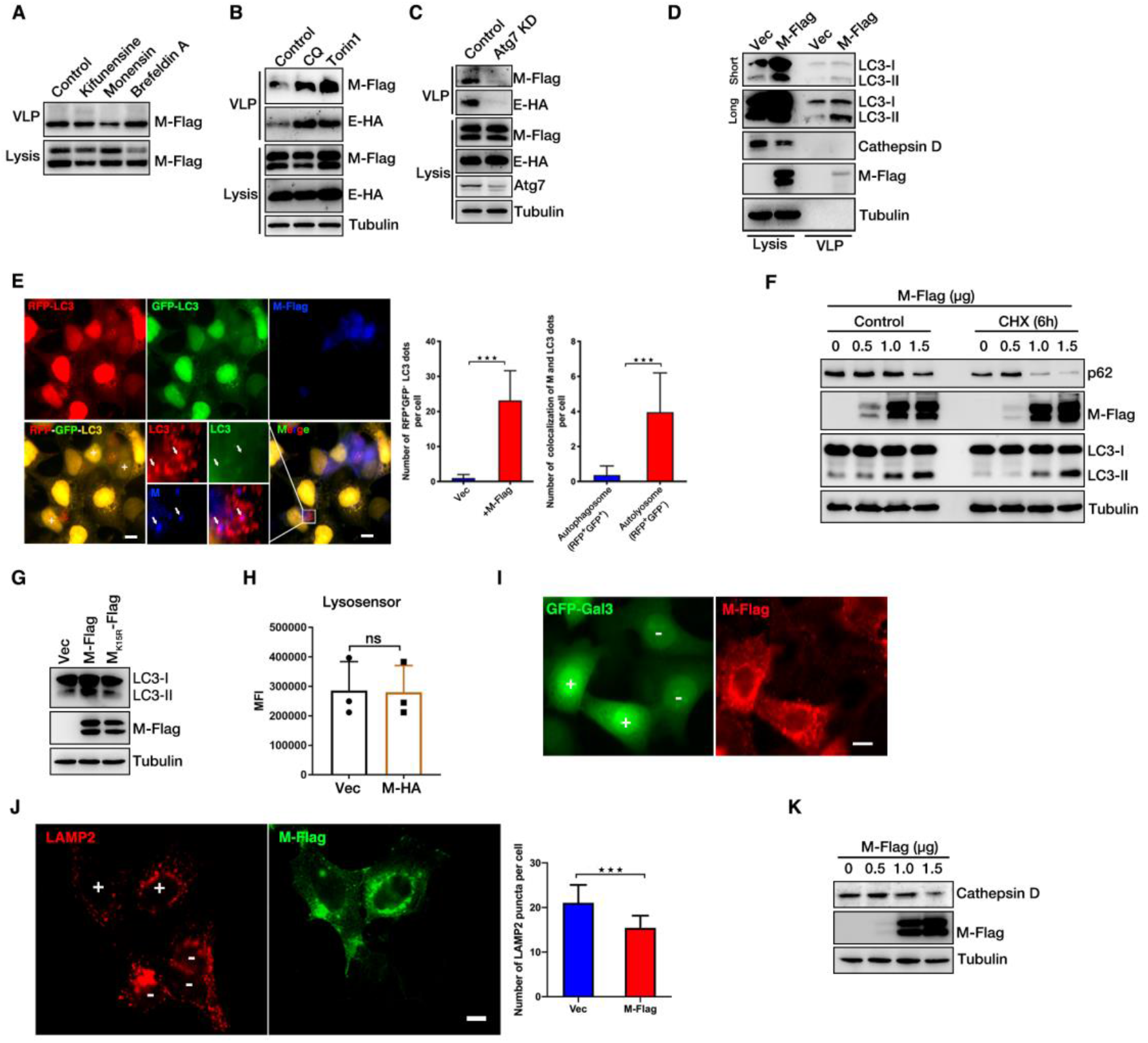
SARS-CoV-2 M Induces Complete Autophagy and Uses Autolysosomes for Virions Release. (A) HEK293T cells transfected with E-HA and M-Flag for 30 h. Cells were further treated with indicated drugs. Lysates and corresponding purified VLPs analyzed via WB. (B) HEK293T cells were transfected with E-HA and M-Flag for 30 h. Cells were further treated with CQ or Torin1 for another 6 h. Lysates and corresponding purified VLPs analyzed via WB. (C) HEK293T cells were transfected with E-HA and M-Flag with or without Atg7 siRNA for 36 h. Lysates and corresponding purified VLPs analyzed via WB. (D) HEK293T cells were transfected with E-HA and M-Flag for 36 h. Lysates were subjected to Flag IP and corresponding purified VLPs analyzed via WB. (E) HeLa cells were transfected with RFP-GFP-LC3, M-Flag and E-HA for 24 h, and cells were analyzed via immunofluorescence. Flag antibody was used for tracking M. Scale bar: 10 μm. The graphs show the quantification of number of RFP+GFP-LC3 dots per cell, and number of colocalization of M and indicated LC3 dots per cell by taking the average number of dots in 50 cells (n = average number of dots in 50 cells). Error bars, mean ± SD of three experiments (n = 3). Student’s t test; ***p < 0.001. (F) HEK293T cells were transfected with or without M-Flag and treated with or without CHX for 6 h, and cells lysis were analyzed via WB. (G) HEK293T cells were transfected with or without M-Flag or M_K15R_-Flag for 36 h, and cells lysis were analyzed via WB. (H) HEK293T cells were transfected with or without M-Flag, and cells were stained by LysoSensor for another 30 min and analyzed via Flow Cytometry. (I) HeLa cells were transfected with GFP-Gal3 and M-Flag for 24 h, and cells were analyzed via immunofluorescence. Flag antibody was used for tracking M. Scale bar: 10 μm. (J) HeLa cells were transfected with M-Flag for 24 h, and cells were analyzed via immunofluorescence. Flag antibody was used for tracking M and LAMP2 antibody was used for tracking lysosomes. Scale bar: 10 μm. The graph shows the quantification of number of colocalization of LAMP2 puncta per cell by taking the average number of dots in 50 cells (n = average number of dots in 50 cells). Error bars, mean ± SD of three experiments (n = 3). Student’s t test; ***p < 0.001. (K) HEK293T cells were transfected with M-Flag for 36 h, and cells lysis were analyzed via WB.

Next, we used the tandem reporter RFP-GFP-LC3 to investigate whether M targets autophagosome or autolysosome. The GFP of this tandem reporter is sensitive and attenuated in an acidic pH environment by lysosomal degradation, whereas the RFP is not. Therefore, the fusion of autophagosomes with lysosomes will result in the loss of yellow fluorescence and the appearance of only red fluorescence. In M and E co-expression cells, many LC3-positive autophagic vacuoles were red (RFP^+^GFP^−^) (**Fig. 5E**). We further found that M targeted red LC3-positive dots (**Fig. 5E**), suggesting that M targets autolysosomes (red LC3 positive dots).

Next, we sought to determine whether M induces autophagy for virion release. For this purpose, we gradually increased the expression of M. To our surprise, we found that LC3-II levels were notably increased in M-expressed cells, whereas we did not observe significant degradation of p62 (**Fig. 5F**). Under CHX treatment, protein translations were inhibited and we then observed the significant increased LC3-II level and degradation of p62 in M-expressed cells (**Fig. 5F**), suggesting that M induces autophagy and autophagic degradation was inhibited. Furthermore, ubiquitination defect mutant K15R failed to induce autophagy (**Fig. 5G**). Thus, these results indicated that M expression induces autophagy which is dependent on RNF5-mediated ubiquitin modification.

Next, we sought to determine why M can use autolysosomes for release, not for degradation. We did not observe any change of lysosome pH (**Fig. 5H**). Galectin3 is specifically localized on damaged endosomes or lysosomes (38). Similar with control cells, we found that GFP-Gal was also diffusely localized in M expressed cells (**Fig. 5I**), suggesting that M does not cause lysosome damage. We further used LAMP2 to track lysosomes and found that the number of LAMP2 positive puncta in M-expressed cells was slightly decreased compared with control cells (**Fig. 5J**). Furthermore, we observed the significant decrease in maturated Cathepsin D level in M-expressed cells compared to control cells (**Fig. 5K**), suggesting that M expression inhibits the degraded activity of lysosome by downregulating the maturation of protease Cathepsin D. Taken together, our data showed that RNF5-mediated ubiquitin modification in M is critical for SARS-CoV-2 virion release through inducing autophagy and using autolysosome.

## Discussion

In this study, we used VLP systems to investigate the molecular mechanisms of SARS-CoV-2 release in detail. We showed that SARS-CoV-2 E interacts with M to promote M self-interaction and ensures the uniform size of SARS-CoV-2 viral particles. M alone forms homo-oligomerization via its CTD. E binds to the CTD and NTD of M to promote the oligomerization of M. We further identified RNF5 act as the E3 ligase of M respectively to regulate the interaction of M with E via ubiquitin modification of M. RNF5 ubiquitinates M at the residue K15 to enhance the interaction of its NTD with E, thus enhances the stability of M-E complex on membrane and ensures the uniform size of VLPs to promote viral release. Knockdown of RNF5 decreases while overexpression of RNF5 increases SARS-CoV-2 VLPs release. SARS-CoV-2 infection experiments also showed that extracellular viral production (virions released into supernatants) is significantly lower in RNF5 KD cells than wild type cells. We further determine the mechanism of how M mediates virion trafficking to cell membrane for release. We found that CQ or Torin1 treatment significantly enhanced VLPs release and knockdown of Atg7 significantly decreased VLPs release, suggesting a critical role of autophagy in virion release. M induces complete autophagy is dependent on RNF5-mediated ubiquitin modification, and inhibits the activity of lysosome protease to block the degradation of autolysosomes, and thus using autolysosomes for release. Altogether, these data revealed a previously undescribed mechanism that RNF5 mediates ubiquitin modification of SARS-CoV-2 M to stable M-E complex and traffic to autolysosomes for virion release.

M homo-oligomerization has been reported to be critical for viral assembly and release: M proteins of paramyxoviruses can self-associate to form higher-order structures (12), VP40 of Ebola virus can oligomerize into hexamers or octamers in vitro via membrane association (39, 40). Crystal structures, biochemistry, and cellular microscopy showed that VP40 can assemble into different structures that contribute to distinct functions, including viral assembly, release, and transcription via structural transformations (41). Herein, we found that M of SARS-CoV-2 forms homo-oligomerization via its CTD which is critical for VLPs release. Different from most of enveloped viruses that M is necessary and sufficient to mediate VLPs release, previous studies have shown that E/M proteins of MHV or M/E/N proteins of SARS-CoV are all required for efficient assembly, trafficking and VLPs release (7). Unlike SARS-CoV, here we found that M and E of SARS-CoV-2 are necessary and sufficient to mediate VLPs release while N is not critical for this process. E interacts with M and enhances the M self-interaction to promote viral release, and E homo-oligomerization is not required for VLPs release. The diameter of SARS-CoV-2 is around 65-125 nm. Viruses use strategies to avoid production of defect virion, whereas the mechanism(s) by how virus could ensure the uniform size of viral particles distribution which is critical for viral maturation were poorly known. M self-interaction is mediated by its CTD bur not NTD, while E binds to both NTD and CTD of M. We speculated that E binds to NTD of M and mediates the self-interaction of NTD of M, and the interaction of E and NTD of M enhances the self-interaction of M to promote the stability of M on membrane with high surface tension and ensure the uniform size of VLPs. Further *in vitro* experiments and structures analysis of M-E complex need to be done to elucidate details by how virus could ensure the uniform size of viral particles distribution.

Different from previous reported function in protein degradation, instead of regulating the stability of M, RNF5 mediated-ubiquitination of M enhances the interaction of NTD of M with E to promote the steady of M-E complex. Notably, a mutant M_K15R_ that is deficient in ubiquitin modification lost its ability for VLPs release, suggesting a critical role of RNF5 in controlling the release and maturation of SARS-CoV-2. Similar with our finding of RNF5 regulates proteins interaction via ubiquitin modification, RNF5 also interacts with JAMP resulting in the Lys-63 chain ubiquitination and such ubiquitination does not alter JAMP stability, but rather decreases its association with proteasome subunits and p97, a key component of the ERAD response (29).

Autophagy is a multistep process by which cytoplasm components were engulfed into autophagosome and shuttled to lysosomes for degradation (42). Autophagy is activated by viral infection and can serve as innate immunity and adaptive immune response against intracellular viruses (43). In addition, viruses have developed strategies to subvert or directly use autophagy for their own production. For example, our previous study showed that phosphoprotein of HPIV3 blocks autophagosome-lysosome fusion to increase virus production (36). ORF3a of SARS-CoV-2 blocks HOPS complex-mediated assembly of the SNARE complex required for autolysosome formation (44). Recent study found that SARS-CoV-2 uses lysosomes for egress instead of the biosynthetic secretory pathway and highlighted the critical role of lysosomal exocytosis for virus egress (37). However, whether autolysosomes could serve as the cargo for SARS-CoV-2 release has not been determined. Here we found that M targets autolysosomes with co-expression of E, and LC3 is co-released with M-mediated VLPs. RNF5 mediated-ubiquitin modification is critical for the autophagy induction of M. Further experiments need to be done to elucidate details by how M induces autophagy and how autolysosomes promote M release. Our unpublished data suggested that M had no interaction with autophagic proteins, suggesting M induces autophagy via a new autophagic regulatory factor. By using siRNA screening, we have identified a novel candidate regulates autophagy and interacts with M which may serve as the key adaptor between autophagy and M.

Combined with previous studies, we and other researchers determine the critical roles of RNF5 in virus infection: RNF5 ubiquitinates STING and MAVS for degradation to decrease the IFNβ response; RNF5 ubiquitinates SARS-CoV-2 M to facilitate virions release. All in all, our findings provide RNF5 as a potential target for antiviral drug development.

## Materials and Methods

### Cell Cultures

HEK293T, *rnf5* KO HEK293T, AD293, HeLa and Vero cells were cultured in Dulbecco’s modified Eagle’s medium (DMEM; GIBCO) supplemented with 10% fetal bovine serum (FBS, Sigma-Aldrich) at 37°C with 5% CO_2_.

### Plasmids Construction

PCDNA4.0-M-Flag, PCDNA4.0-M_Δ_TMD1-Flag, PCDNA4.0-M_Δ_TMD2-Flag, PCDNA4.0-M_Δ_TMD3-Flag, PCDNA4.0-M_Δ_NTD-Flag, PCDNA4.0-M_Δ_40-50aa-Flag, PCDNA4.0-M_Δ_74-77aa-Flag, PCDNA4.0-M_Δ_CTD-Flag, PCDNA4.0-M_Δ_20-100aa-Flag, PCDNA4.0-M-CTD-Flag, PCDNA4.0-M-20-100aa-Flag, PCDNA4.0-M-K14R-Flag, PCDNA4.0-M-K15R-Flag, PCDNA4.0-M-K50R-Flag, PCDNA4.0-M-K162R-Flag, PCDNA4.0-M-K166R-Flag, PCDNA4.0-M-K180R-Flag, PCDNA4.0-M-K205R-Flag were cloned into mammalian expression vector PCDNA4.0-Flag. PCDNA4.0-M-HA, PCDNA4.0-E-HA, PCDNA4.0-RNF5-HA, PCDNA4.0-RNF5_Δ_TMD-HA and PCDNA4.0-RNF5-C42S-HA were cloned into mammalian expression vector PCDNA4.0-HA. PCDNA3.0-HA-UB was cloned into mammalian expression vector PCDNA3.0-HA. PCDNA4.0-E-Myc and PCDNA4.0-DFCP1-Myc were cloned into mammalian expression vector PCDNA4.0-Myc. PTY-M-HA was cloned into mammalian expression vector PTY-HA. PCDNA4.0-EGFP-E-HA, PCDNA4.0-EGFP-E_Δ_NTD-HA and PCDNA4.0-EGFP-E_Δ_CTD-HA were cloned into mammalian expression vector PCDNA4.0-EGFP-HA.

### Oligonucleotides

RNF5, GPAT4, ACSL3, PGAM5, AGPAT4, AUP1, DERL1, AGPAT5, SEL1L, OS9, FAM8A1, DERL2, UBE2J1, UBE2G2, RIINT1, RNF126, RNF138, RNF170, POH1, ZW10 and EDEM3 siRNAs were purchased from Guangzhou RiboBio. pSuper RNF5 shRNA AGCTGGGATCAGCAGAGAG.

### Antibodies and Reagents

Mouse monoclonal anti-FLAG (F1804), anti-HA (H3663) and ERGIC53 (SAB2101351) were obtained from Sigma-Aldrich. Anti-Myc (2278) was obtained from Cell Signaling Technology. Anti-RNF5 (sc-81716), LAMP1 (sc-65236) and LAMP2 (sc-18822) were obtained from Santa Cruz Biotechnology. Rabbit anti-LC3 (PM036) and mouse anti-LC3 (M152-3) were obtained from MBL. Anti-Tubulin (E7-S) was obtained from Developmental Studies Hybridoma Bank. Mouse anti-p62 (H00008878-M01) was obtained from ABnOVA. Goat anti-Mouse IgG (H+L) Secondary Antibody, Alexa Fluor® 568 conjugate (A11031), goat anti-Rabbit IgG (H+L) Secondary Antibody, Alexa Fluor® 568 conjugate (A11036), goat anti-Mouse IgG (H+L) Secondary Antibody, Alexa Fluor® 488 conjugate (A32723), goat anti-Rabbit IgG (H+L) Secondary Antibody, Alexa Fluor® 488 conjugate (A32731) and goat anti-Mouse IgG (H+L) Cross-Adsorbed Secondary Antibody, Alexa Fluor® 405 (A31553) were obtained from Thermo Fisher Scientific. Anti-Myc magnetic beads (B26301) and Anti-HA magnetic beads (B26201) were obtained from Bimake. Anti-Flag M2 Affinity Gel (A2220) was obtained from Sigma-Aldrich. Monensin Solution (50501ES03) was obtained from YEASEN Biotech. Kifunensine (GC17735) was obtained from Glpbio. Brefeldin A (B5936) was obtained from Sigma-Aldrich. Torin 1 (F6101) was obtained from Ubiquitin-Proteasome Biotechnologies.

### Immunoprecipitation and Western Blot

Cells were harvested and lysed with TAP buffer (20 mM Tris-HCl, pH 7.5, 150 mM NaCl, 0.5% NP-40, 1 mM NaF, 1 mM Na3VO4, 1 mM EDTA, Protease cocktail) for 30 min on ice. The supernatants were collected by centrifugation at 13000 rpm for 20 min at 4°C. For Flag, Myc or HA tag IP, tag affinity gel beads were added in supernatants and incubated overnight. Beads were washed three times with TAP buffer and boiled at 100°C for 10 min in SDS protein loading buffers and analyzed by WB. Protein concentration was determined based on the Bradford method using the Bio-Rad protein assay kit. Equal amounts of protein were separated by 12% SDS-PAGE and electrophoretically transferred onto a nitrocellulose membrane. After blocking with 5% nonfat milk in PBST, membrane was incubated with the primary antibodies, followed by HRP-conjugated goat anti-mouse IgG.

### VLP release assay

To analyze the VLPs released from cells, the culture medium of transfected cells was collected and centrifuged at 13,000 rpm for 5 min to remove cell debris, then layered onto a cushion of 20% (wt/vol) sucrose in PBS, and subsequently ultracentrifuged on OptimaTM MAX-XP Ultracentrifuge(BECKMAN) at 35,000 rpm for 2 hr at 4°C; the VLPs pelleted at the bottom of the tubes were resuspended in 35 ul of TNE buffer (50 mM Tris-HCl [pH 7.4], 100 mM NaCl, 0.5 mM EDTA [pH 8.0]) overnight at 4°C. Samples were boiled with SDS-PAGE loading buffer and analyzed by western blotting as described above.

### Protease protection assay

VLPs from medium of cells were prepared as described above. Trypsin (GIBCO) was added to the final concentration at 2 μg/ml, along with 1% Triton X-100 if desired. Samples were incubated at 37°C for 1 h, then mixed with SDS-PAGE loading buffer, and boiled for WB analysis.

### SARS-CoV-2 Virus infection

All works with live SARS-CoV-2 virus were performed inside biosafety cabinets in the biosafety level 3 facility at Hubei Provincial Center for Disease Control and Prevention. Vero cells in 6-well plates were infected with SARS-CoV-2 (WBP-1) at a MOI of 1 PFU/cell for 1 h at 37°C with 5% CO_2_, then infection medium was removed and replaced with fresh DMEM medium with 2% FBS.

### Plaque Assay

SARS-CoV-2-containing culture medium was serially 10-fold diluted. Vero cells in 6-well plates were grown to 60 to 70% confluency and infected with 100 μl of the dilutions. Plates were incubated for 2 h at 37°C with 5% CO_2_, and then washed with PBS, the infection medium was replaced with methylcellulose, and plates were incubated at 37°C with 5% CO_2_ for another 3 to 4 days until visible viral plaques were detected. Plates were stained with 0.5% crystal violet for 4 h at room temperature and washed; then the plaques were counted and the viral titers were calculated.

### Immunofluorescence Analysis

Cells were washed with PBS and fixed with 4% paraformaldehyde for 15 min in room temperature and then cells were washed three times with PBS and then incubated with 0.1% saponin for 10 min. After washing three times with PBS, cells were blocked with 10% FBS for 30 min. Specific primary Abs were added and incubated overnight, and cells were then washed with PBS for three times, followed by incubation with the goat anti-rabbit IgG Rhodamine or goat anti-mouse IgG fluorescein secondary antibody for 1 h. Cells were then washed with PBS for three times.

### Quantification and statistical analysis

Statistical parameters including the definition and exact values of n, distribution and deviation are reported in the figure legends. Data are expressed as mean ± standard deviation (SD). The significance of the variability between different groups was determined by two-way analyses of variance using GraphPad Prism software. Error bars, mean ± SD of two or three independent experiments. Student’s t test, a p value of < 0.05 was considered statistically significant and a p value of > 0.05 was considered statistically non-significant (NS).

## Acknowledgments

We thank Dr. Nevan J. Krogan (UCSF) for providing SARS-CoV-2 viral proteins expression plasmids; Dr. Qiyun Zhu (Lanzhou Veterinary Research Institute) and Yingjie Sun (Shanghai Veterinary Research Institute, CAAS) for providing *rnf5* KO cells; Dr. Xiaochen Wang (Chinese Academy of Science) for GFP-Gal3; Dr. Anbing Shi (HUST) for helpful discussion. This work was supported by the Major Research Plan of the National Natural Science Foundation of China (92054107, B.D.); National Natural Science Foundation of China (82041004); Start-up funds from Huazhong University of Science and Technology (3011510035, B.D.); HUST COVID-19 Rapid Response Call (2020kfyXGYJ036); Zhejiang University special scientific research fund for COVID-19 prevention and control (2020XGZX089).

## Author contributions

Z.Y. performed most of the experiments; Y.W. contributed with autophagy and lysosome experiments; X.T. and H.X. help with repeating experiments; H.X. and M.Y. contributed with constructs; K.C. and B.H. contributed with SARS-CoV-2 infection experiments. Y.L. contributed with materials; K.T. contributed with image and NanoSight NS300 (Malvern) assay and edited the manuscript; B.D. conceived the project, designed the experiments, analyzed the data and wrote the manuscript. All authors discussed the results and commented on the manuscript.

## Data and materials availability

All data needed to evaluate the conclusions in the paper are present in the paper and/or the Supplementary Materials. Additional data related to this paper may be requested from corresponding author.

