## supplement for "The E3 Ubiquitin Ligase RNF5 Facilitates SARS-CoV-2 Membrane Protein-Mediated Virion Release"

### Figures

#### Fig S1. SARS-CoV-2 M Interacts with RNF5 to Mediate Viral Release.

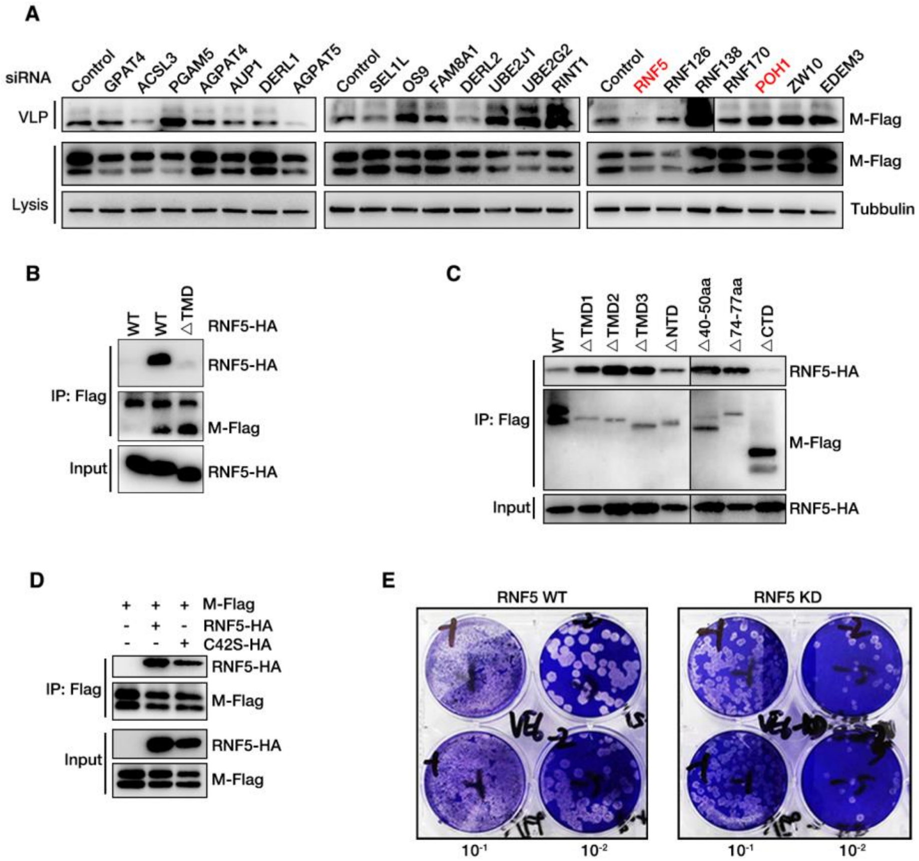


(A) HEK293T cells transfected with indicated siRNAs and further transfected with E-HA and M-Flag for 36 h, then cell lysates and corresponding purified VLPs were analyzed via WB. (B) HEK293T cells were transfected with M-Flag and RNF5-HA or its mutant for 36 h, and subjected to Flag IP and analyzed via WB. (C) HEK293T cells were transfected with RNF5-HA and M-Flag or its mutant for 36 h, and subjected to Flag IP and analyzed via WB. (D) HEK293T cells were transfected with indicated plasmids for 36 h, and subjected to Flag IP and analyzed via WB.

(D) RNF5 WT and KD Vero-E6 cells were infected with SARS-CoV-2 (WBP-1) for 24 h and then mediums were collected and analyzed via Plaque Assay.
